# Dissociating decisional and temporal information in interval categorisation

**DOI:** 10.1101/659375

**Authors:** Vanessa C. Morita, João R. Sato, Marcelo S. Caetano, André M. Cravo

## Abstract

Interval timing is fundamental for humans and non-human animals to interact with their environment. Several studies that investigate temporal processing combine behavioural tasks with neurophysiological methods, such as electrophysiological recordings (EEG). However, in the majority of these studies, it is hard to dissociate whether EEG activity reflects temporal or decisional information. In the present study, we investigated how time and decision is encoded in the EEG signal while human participants performed a temporal categorisation task with two different temporal references. Using a combination of evoked potentials and multivariate pattern analysis, we show that: (1) During the interval to-be-timed, both temporal and decisional information are encoded; (2) Activity evoked by the end of the interval encodes almost exclusively decisional information. These results suggest that decisional aspects of the task better explain EEG activity commonly related to temporal processing. The interplay between the encoding of time and decision is consistent with recent proposals that approximate temporal processing with decisional models.

## 1 Introduction

The perception of time in the range of hundreds of milliseconds to seconds is fundamental for humans and non-human animals to interact with their environments [Buhusi and Meck, 2005; Paton and Buonomano, 2018]. Although several theoretical models have been proposed to explain how the brain keeps track of time in this range, there is still no evidence toward a single one [Hass and Durstewitz, 2016]. It is often hard to compare different models due to their mathematical similarity and to their overlap in behavioural predictions [Balci and Simen, 2016]. For this reason, studies that investigate temporal processing in humans often combine behavioural tasks with different neurophysiological methods, such as electrophysiological recordings (EEG). Based on the patterns of EEG activity, one could dissociate between different theoretical models. However, for this strategy to work, it is essential to determine what aspects of the task these EEG patterns are measuring.

A common approach to investigate interval timing is to use temporal categorisation tasks, in which participants judge whether a given interval is longer or shorter than a temporal reference. Different EEG markers have been proposed as indexing temporal information in this kind of task (for a recent review see [Kononowicz et al., 2016]). Perhaps the most famous is the contingent negative variation (CNV), a slow cortical potential of developing negative polarity at frontocentral scalp locations evoked during the to-be-timed interval. It has been consistently found that the CNV peaks at the moment of the reference interval [Macar and Vidal, 2003; Pfeuty et al., 2003; Pouthas et al., 2000; Wiener and Thompson, 2015]. However, recent studies have questioned the role of the CNV in time perception (for a review, see [Kononowicz et al., 2016]) and have suggested that post-interval event-related potentials (ERPs), such as the Late Positive Component of timing (LPCt) [Lindbergh and Kieffaber, 2013; Paul et al., 2011; Wiener and Thompson, 2015] and the N1-P2 complex [Kononowicz and van Rijn, 2014] reflect subjective timing better.

A significant limitation of the majority of temporal categorisation studies is to use a single temporal reference. When this is the case, it is hard to dissociate whether EEG activity reflects temporal or decisional information. For example, when asked to judge whether a test interval of 1100 ms is shorter or longer than a reference of 900 ms, it is not possible to separate whether EEG activity is reflecting the physical duration of the interval or the fact that it is shorter than the reference. One method to differentiate between these possibilities is to use different temporal references in separate blocks of the same task. Consider if, on the above example, the same 1100 ms interval was presented on a different block in which participants had to judge whether it was shorter or longer than a reference of 1500ms. By comparing brain activity elicited by the 1100 ms in the different blocks, it would be possible to dissociate whether this brain activity reflects the duration of the interval or its relation to the temporal reference. In one of the few studies that used this approach in humans, Pfeuty and colleagues found that the CNV peaked at the moment of the reference interval and that its slope was steeper for shorter references Pfeuty et al. [2005]. However, there is still a shortage of studies that have used this approach and an even smaller number of studies that have used it to investigate post-interval event-related potentials.

In the present study, we investigated how temporal and decisional information is encoded in the EEG signal while participants performed a temporal categorisation task with two different temporal references. We combined ERP and multivariate pattern analysis (MVPA) to analyse time-resolved EEG signals. Our results suggest that during the interval to-be-timed, both temporal and decisional information is encoded. On the other hand, the post-interval signal encodes more strongly decisional information.

## 2 Results

### 2.1 Behavioural Results

Participants performed temporal categorisations in a computerised “shoot the target” task. The task consisted of two types of trials. In regular trials (figure 1A), a visual target transited from the left periphery of the screen towards a central target zone, where there was an “aiming sight”. In these trials, participants were instructed to press a button to produce a “shot” (an audiovisual stimulus) when the target passed the aiming sight. In test trials (figure 1B), the trajectory of the target was masked by an occluder, and only the aiming sight remained visible throughout the entire trial. In these trials, participants did not see the movement of the target, but heard the start of its movement by a characteristic sound that targets made when they started moving (both in regular and in test trials). Critically, in test trials, participants were instructed that they did not have to produce the shot themselves and that one automatic shot would be presented after a certain interval. Participants had to judge whether the shot was given “before” or “after” the target reached the centre of the screen (see methods for more detail).

**Figure 1:**
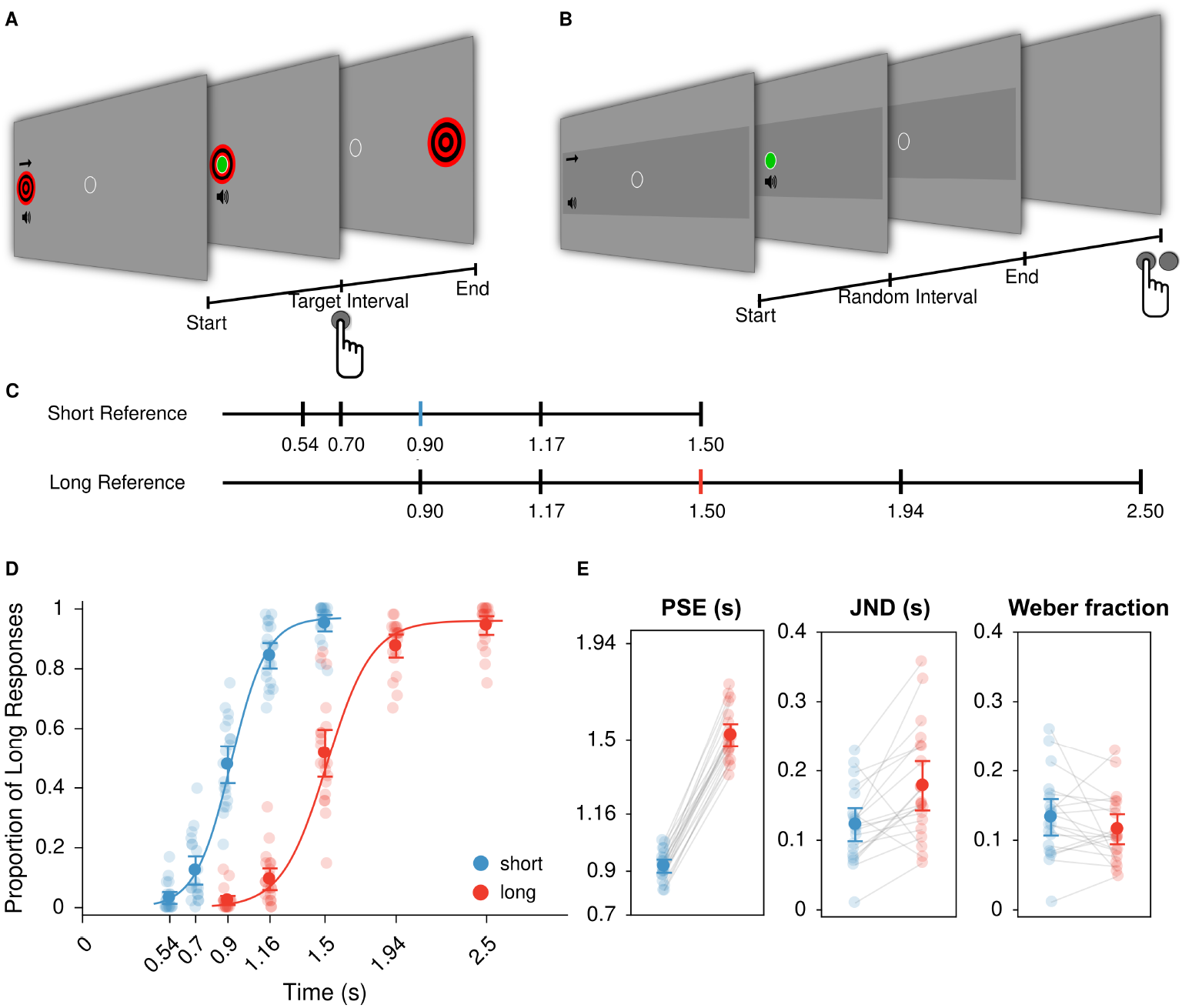
Experimental task and behavioural results. (A) The task consisted of a computerised “shoot the target” task. In regular trials, participants had to shoot a target at the reference interval (0.9 or 1.50 seconds). (B) In test trials, one of five possible intervals was presented. Participants had to judge whether the shot occurred after an interval “shorter” or “longer” than the respective reference interval. (C) Schematic representation of test intervals for each reference interval. (D) Psychometric functions adjusted to the proportion of “after” responses (mean ± confidence interval) for the short and long reference intervals (blue and red, respectively). Lighter dots indicate data from each participant. (E) Estimated parameters: Point of Subjective Equality (PSE), Just Noticeable Difference (JND) and Weber fraction (mean ± confidence interval). Gray lines connect data from the same participant.

In different blocks, the target could move at two different speeds (figure 1C): in short blocks, the target reached the aiming sight after 0.9 seconds. In long blocks, it took 1.5 seconds to reach the aiming sight. The intervals in which the shot could be presented in test trials were also block-dependent. In short blocks, the shot could be presented in one of the following five intervals: 0.54 s, 0.70 s, 0.90 s, 1.16 s or 1.5 s. In long blocks, the intervals were of 0.90 s, 1.16 s, 1.5 s, 1.94 s or 2.50 s. Critically, the intervals used for each reference were chosen such that the three longest intervals of the short reference were the same as the three shortest intervals of the long reference.

The proportion of “after” responses was calculated for each test interval and reference and a logistic psychometric function was fitted for each condition. Participants made few mistakes at test intervals which were clearly shorter or longer than the reference intervals (figure 1D). The estimated Point of Subjective Equality (PSE) was of 0.92 ± 0.02 s (mean ± s.e.m., 95% Confidence Interval [CI] = 0.89, 0.95) for the short reference interval and 1.52 ± 0.03 s (CI = 1.47, 1.57) for the long reference interval, with extreme evidence that they were different (paired t-test *t*_19_ = −29.6474, *p* < 0.001, *d* = −6.6294, *BF*_10_ = 1.6685 × 10^14^).

Temporal sensitivity for each reference interval was assessed by calculating the Just Noticeable Differences (JND, figure 1E). We found strong evidence that the JND was smaller for the short reference interval (JND_*short*_ = 0.12 ± 0.01s, CI = 0.09, 0.15, JND_*long*_ = 0.18 ± 0.02s, CI = 0.14, 0.21, paired t-test *t*_19_ = −3.6281, *p* = 0.0018, *d* = −0.8113, *BF*_10_ = 22.0787), suggesting that shorter intervals were easier to discriminate than the longer ones. A key aspect of time perception is the scalar property, a particular case of the Weber Law, which states that the variability on time estimation increases proportionally to the magnitude of the interval. We calculated the Weber fractions and found anecdotal evidence in favour that the Weber fractions for short and long reference intervals are similar (Weber_*short*_ = 0.13 ± 0.01, CI = 0.10, 0.16, Weber_*long*_ = 0.12 ± 0.01, CI = 0.09, 0.14, paired t-test, *t*_19_ = 1.3258, *p* = 0.2, *d* = 0.2964, *BF*_10_ = 0.4977).

### 2.2 Pre-interval EEG activity

#### 2.2.1 CNV analysis

In a first step, we analysed the contingent negative variation (CNV), an ERP commonly associated with temporal processing. Figure 2 shows the evoked potentials (grand mean across trials and participants) at central electrodes for each test interval and reference. As can be seen, the CNV had an increasing negative amplitude, peaked at the reference interval and then had its amplitude decreased.

**Figure 2:**
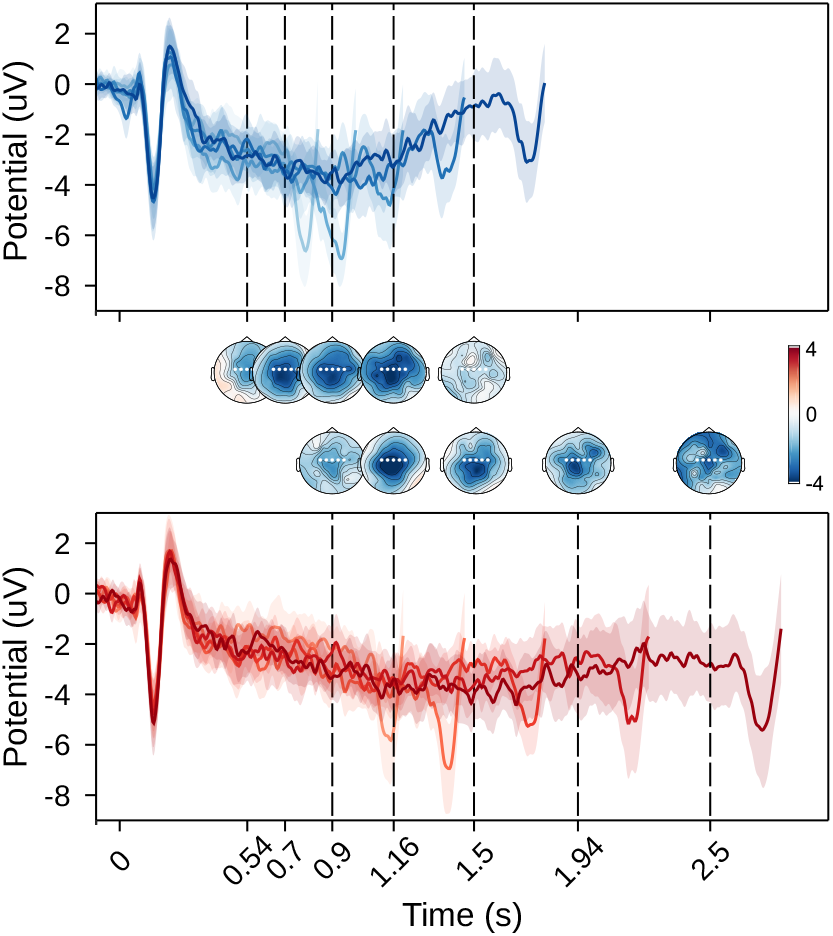
Contingent Negative Variation. Grand-average ERPs for each test interval (darker colours corresponding to longer intervals, mean ± confidence interval) and reference (blue for the short reference and red for the long reference), estimated at central electrodes. Zero indicates the onset of the interval, while dashed lines indicate the offset of each test interval. Topographies shows the mean potential in a window of 100ms before the presentation of the shot for each test interval.

We adapted an approach used by Pfeuty et al. [2005] to be able to estimate, simultaneously, the moment in which the CNV peaked, its maximum amplitude, and its slope. This was done by adjusting two linear functions with a single but inverted slope to the longest test interval of each condition (figure 3). We found extreme evidence that the CNV peaked earlier in blocks in which the temporal reference was shorter (Peak_*short*_ = 0.88 ± 0.05 s, CI = 0.78, 0.98, Peak_*long*_ = 1.37 ± 0.12 s, CI = 1.13, 1.61, paired t-test, *t*_19_ = −4.8105, *p* < 0.001, *d* = −1.0757, *BF*_10_ = 233.8504). We further compared whether the CNV peaked at the moment of the temporal reference (0.9s or 1.5s). For the short reference, we found moderate evidence that the CNV peaked at the moment of the temporal reference (one-sample t-test against 0.9s, *t*_19_ = −0.2941, *p* = 0.7719, *d* = −0.0658, *BF*_10_ = 0.2416) and anecdotal evidence that this was the case for the long reference (one-sample t-test against 1.5s, *t*_19_ = −1.0457, *p* = 0.3088, *d* = −0.2338, *BF*_10_ = 0.3761).

**Figure 3:**
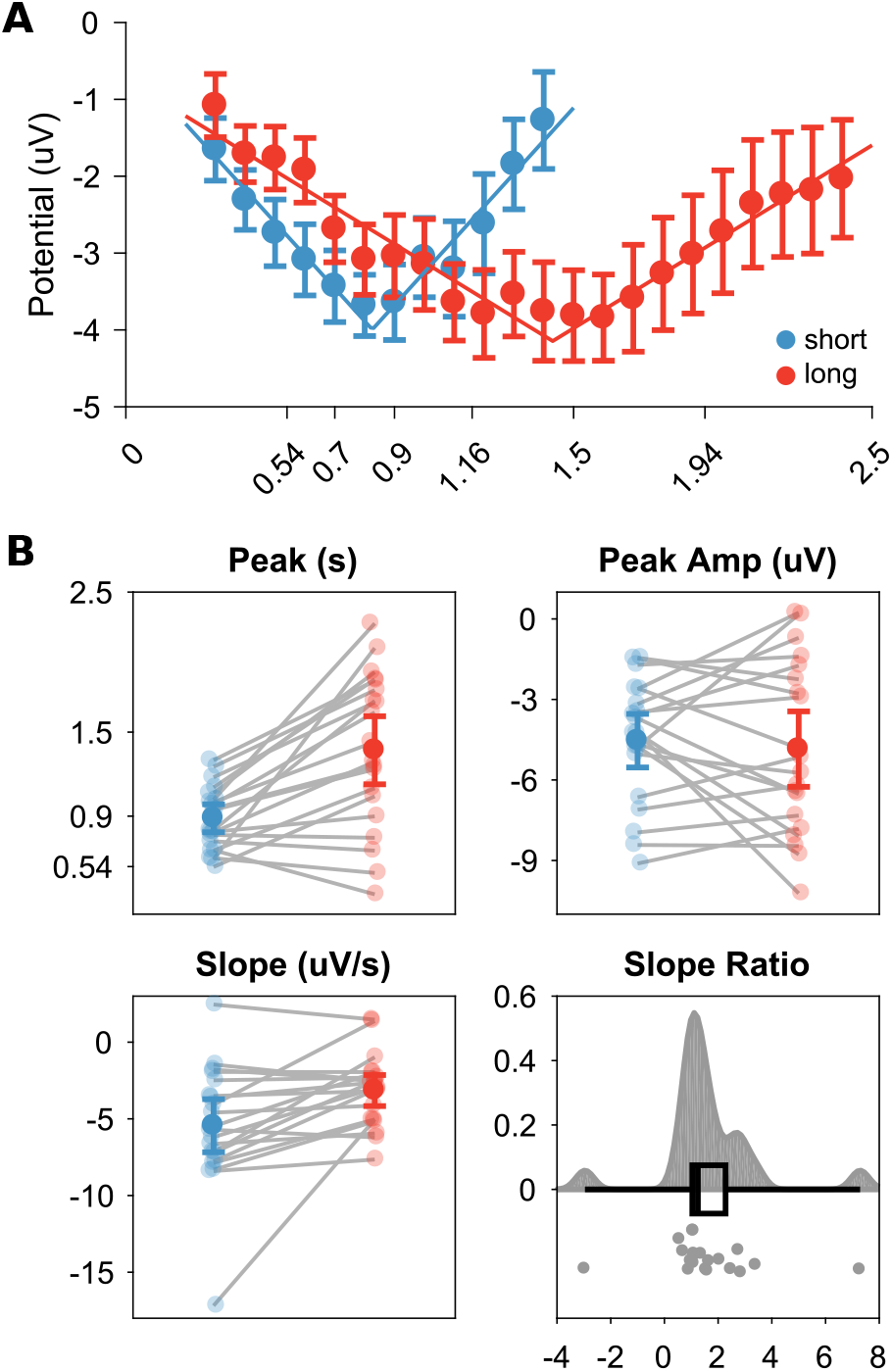
CNV dynamics for different temporal references. (A) Grand-averaged ERPs for the longest interval of each reference (short reference in blue and long reference in red) binned in 100ms windows. Lines shows the fit of the linear model. (B) Parameters fitted in the model (the moment in which the CNV peaked, its amplitude at the peak, its slope and the ratio between estimated slopes Slope_*short*_/Slope_*long*_).

In a previous study, Pfeuty et al. [2005] found that the slope of the CNV was steeper when the temporal reference was shorter. In our results, we found moderate evidence of a steeper CNV for the short reference (Slope_*short*_ = −5.45 ± 0.88 *μ*Vs, CI = −7.17, −3.72, Slope_*long*_ = −3.16 ± 0.52 *μ*Vs, CI = −4.17, −2.14, paired t-test, *t*_19_ = −3.1602, *p* = 0.0052, *d* = −0.7067, *BF*_10_ = 8.9414). As slopes were different between conditions, we compared whether the ratio between slopes (Slope_*short*_/Slope_*long*_ = 1.64 ± 0.41) was equal to the ratio between the temporal references (Reference_*long*_/Reference_*short*_=1.66). We found moderate evidence that this was the case (*t*_19_ = −0.0611, *p* = 0.9519, *d* = −0.0137, *BF*_10_ = 0.2327).

Lastly, we compared what was the amplitude of the CNV at its peak and compared this amplitude between conditions. We found moderate evidence that they were similar (Amplitude_*short*_ = −4.53 ± 0.51*μ*V, CI = −5.53, −3.53, Amplitude_*long*_ = −4.85 ± 0.72*μ*V, CI = −6.25, −3.44, paired t-test, *t*_19_ = 0.5356, *p* = 0.5984, *d* = 0.1198, *BF*_10_ = 0.2643).

#### 2.2.2 MVPA

For the pre-interval activity, we focused the analysis in the last moments of each interval. We used a representational similarity analysis (RSA) approach [Kriegeskorte and Kievit, 2013] to investigate how temporal and decisional information influenced neural coding. The logic of this approach was to examine whether distance in time or decision between two conditions led to more distinct EEG patterns. The empirical dissimilarity matrix, which measured how different two conditions were, was built performing pairwise comparisons between the intervals using an l2-regularised logistic classification. The accuracy for each comparison was transformed into d-prime.

To investigate the representation of these different aspects of the task in EEG activity, two theoretical matrices of pairwise dissimilarity between intervals were built: the first based on how distant intervals were on time (physical difference between intervals, 4A, left panel) and the second based on how distant they were on the decision dimension (distance from the reference interval, 4A, right panel).For ease of comparison, all distances were scaled between 0 and 1.

As can be seen in Figure 4A, the upper-left and lower-right quadrants of these theoretical matrices show comparisons within each temporal reference while the upper-right and lower-left quadrant (which are symmetrical) show comparisons of intervals between temporal references. Notice that for *within* reference comparisons (upper-left and lower-right quadrants) there is a strong correlation between the two theoretical matrices. This correlation is expected, given that when a single temporal reference is used, there is a confusion of how distant two moments are in time and how different they are from the reference.

**Figure 4:**
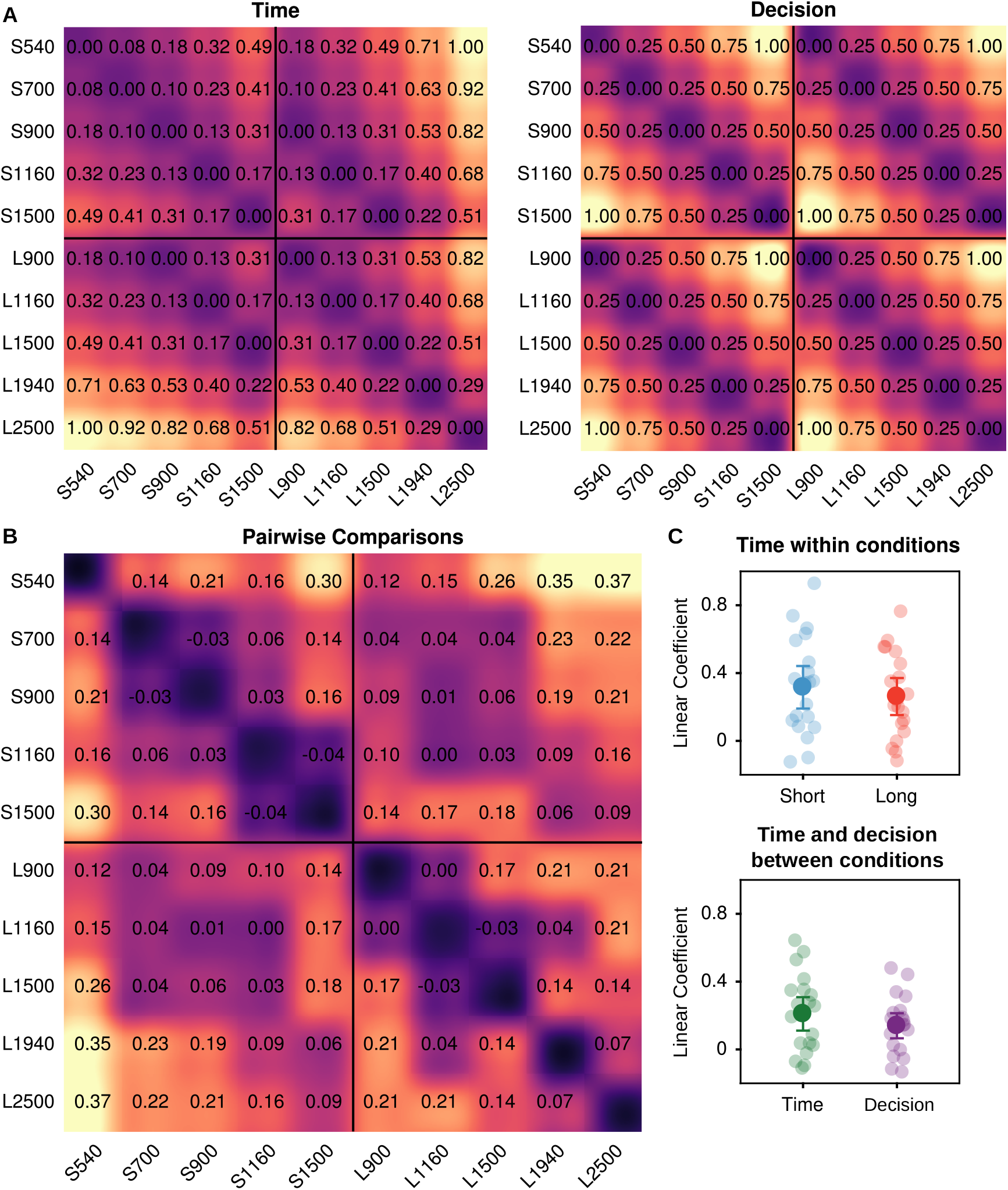
Multivariate analysis for pre-offset EEG signals. (A) Theoretical matrix of discriminabilities between test intervals. Left panel shows the expected discriminability between intervals if EEG data correlates only with time information. Right panel shows the expected discriminability if EEG data correlates only with distance from the reference interval. (B) Experimental matrix of discriminability between intervals. Each value indicates the mean d-prime for the pairwise logistic regression fitted to pre-offset EEG activity (mean of the 100 ms before the offset). (C) Estimated linear coefficients. Top panel shows the linear coefficients (mean ± confidence interval) for each condition using the theoretical matrix for time as a predictor. Bottom panel shows the linear coefficients of a multiple regression using both time and decision theoretical matrices as predictors.

On the other hand, the correlation between the two theoretical matrices is lower for between reference comparisons (upper-right and lower-left quadrants), given that, in these comparisons, different decisions are expected for the same physical interval. For these comparisons, at least two possible outcomes can be expected. If pre-interval EEG activity carries information only about physical time, then just how distant the intervals are in time should modulate their discriminability. For example, the 900 ms interval for the short reference should be indistinguishable from the 900 ms interval for the long reference. If, however, the EEG signal carries information about a decision, i.e., how different each interval is from the temporal reference, then discriminability should depend not on time itself, but the distance from the reference. In this case, for example, the 900 ms interval in the short condition should be more similar to the 1500 ms interval of the long condition.

For the within reference comparison, a linear regression between the theoretical distance in time and the observed discriminability was performed (Figure 4B and Figure 4C). There was extreme evidence of association between distance in time and the experimental dissimilarity for both short and long reference intervals (linear coefficient for short condition: 0.32 ± 0.06, CI = 0.19, 0.44, one-sample t-test against zero, *t*_19_ = 4.9378, *p* < 0.001, *d* = 1.1041, *BF*_10_ = 301.6033; for long condition: 0.26 ± 0.05, CI = 0.15, 0.37, one-sample t-test *t*_19_ = 4.6943, *p* < 0.001, *d* = 1.0497, *BF*_10_ = 185.3083).

For the between references comparisons, we performed a multiple linear regression for discriminability scores with distances in time and in decision as predictors (Figure 4C, lower panel). We found strong evidence for a association for both predictors (linear coefficient for time: 0.21 ±0.05, CI = 0.11, 0.31, one-sample t-test *t*_19_ = 4.1602, *p* < 0.001, *d* = 0.9302, *BF*_10_ = 63.5201; for reference: 0.14 ± 0.04, CI = 0.07, 0.21, one-sample t-test *t*_19_ = 3.6982, *p* = 0.0015, *d* = 0.8269, *BF*_10_ = 25.3441).

### 2.3 Post-interval EEG activity

We used a similar MVPA analysis for post-interval EEG activity (i.e. activity evoked by the shot, indicating the end of the to-be-judged interval) for each time-point (figure 5A, left panel). The within reference analysis showed an association between time and discriminability for both short and long temporal references (window tested=-50ms-700ms; short reference cluster=248-688ms, *p* < 0.001; long reference cluster=184-548ms, *p* < 0.001). As can be seen in Figure 5A (right panel), for the post-interval period there was strong to extreme evidence for an association between distance in time and pattern discriminability starting around 200ms after the end of the interval for both short and long temporal references.

**Figure 5:**
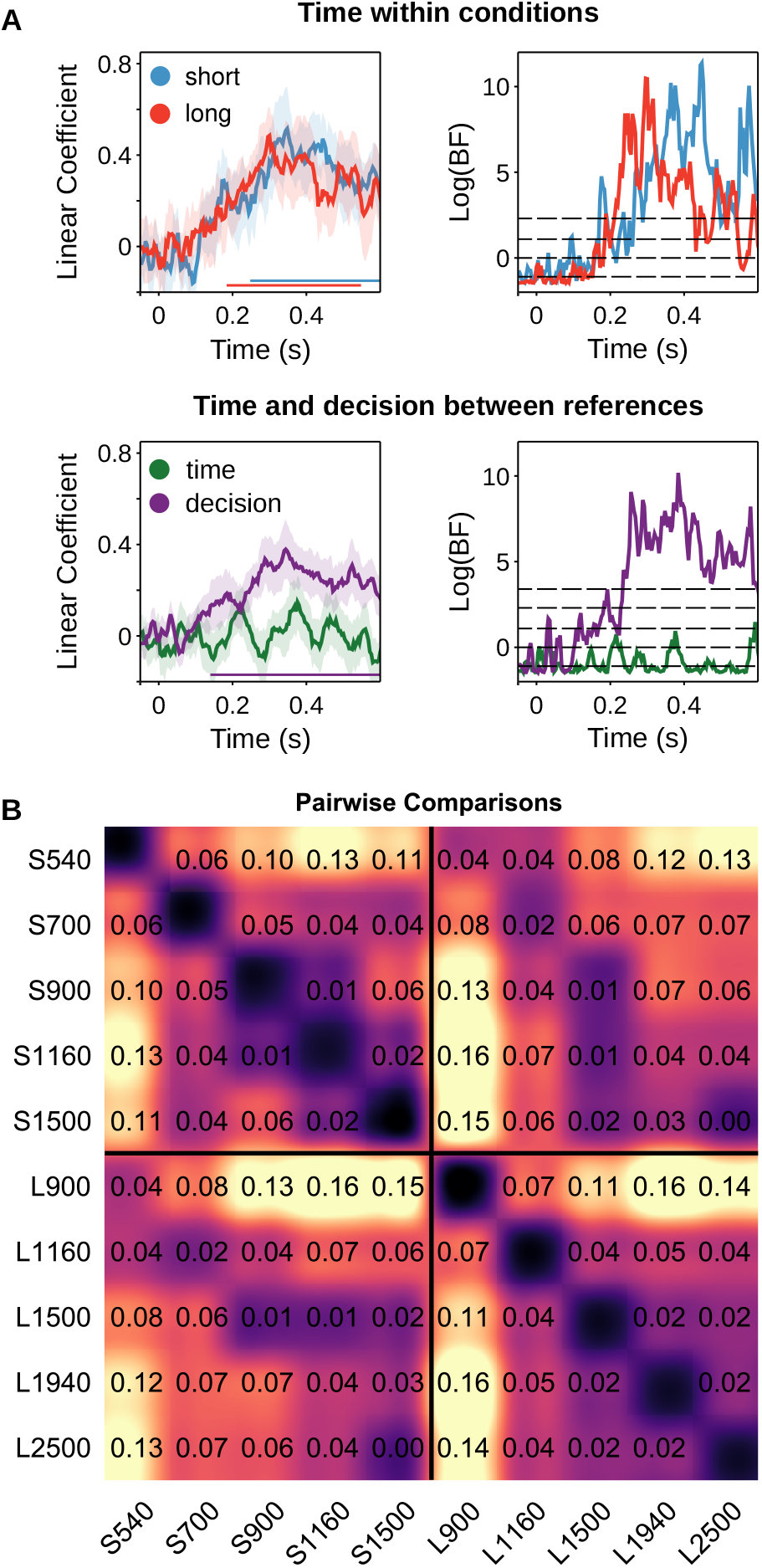
Multivariate analysis for post-offset EEG signals. (A) Upper panels show how distance in time modulated pattern discriminability when comparing intervals within each temporal reference. Left panel shows the estimated coefficients (using distance in time as a predictor) fitted to d-primes for each condition. Bottom coloured lines shows the extension of the temporal cluster. Right panel shows the respective Bayes Factors. For ease of representation Bayes Factors are shown log-transformed. The horizontal dashed lines above zero indicate moderate and strong evidence in favour of the alternative hypothesis, while dashed line below zero indicates level of moderate evidence in favour of the null hypothesis. Lower panels show how distance in time and from the temporal reference modulated pattern discriminability when comparing intervals between each temporal references. Left panel shows the estimated coefficients (both distance in time and of reference as predictors) fitted to d-primes for between conditions pairwise comparisons. Right panel shows the respective Bayes Factors following the same display as above. (B) Mean d-primes for pairwise comparisons between 250 and 550 ms after the offset.

For between references comparisons, there was an association between decisional information and discriminability (window tested=-50ms-700ms; cluster=140-684ms, *p* < 0.001), but not for time (window tested=-50ms-700ms, no candidate cluster). As shown in Figure 5B, on the post-interval period there was strong to extreme evidence for an association between distance in decision and discriminability starting also around 200ms after the end of the interval. For the association between distance in time and discriminability, there was, during the majority of the period post-interval, moderate evidence in favour of no association between them.

To illustrate these results, the dissimilarity matrix for the period between 250 and 550 ms is shown in Figure 5C. As can be seen, during this period, the pattern of discriminability reflects mainly how much shorter or longer a given interval is from its temporal reference. This can be seen, for example, by how the short 900 ms interval is less discriminable from the 1500ms than for the other long intervals.

To compare our findings with previous results that investigate post-interval ERPs and temporal processing, we looked at the pattern of ERPs in two periods: (1) An early frontocentral activity from 100 ms to 200 ms (figure 6A); (2) A later parietal activity from 300 ms to 450 ms (figure 6B). To quantify the size of the effect for time and decision in these periods, we calculated the coefficient of partial determination (CPD) between the amplitude of the ERP with each predictor (interval duration and log-distance from temporal reference). As shown in Figure 6, the amplitude of the ERP was driven mainly by how distant from the reference the interval was for both the early frontocentral activity (CPD_*decision*_ = 0.2768 ± 0.0553, CI = 0.1685, 0.3852; CPD_*time*_ = 0.1491 ± 0.0380, CI = 0.0746, 0.2236) and the late parietal activity (CPD_*decision*_ = 0.2468 ± 0.0517, CI = 0.1455, 0.3482; CPD_*time*_ = 0.1243 ± 0.0399, CI = 0.046, 0.2025).

**Figure 6.**
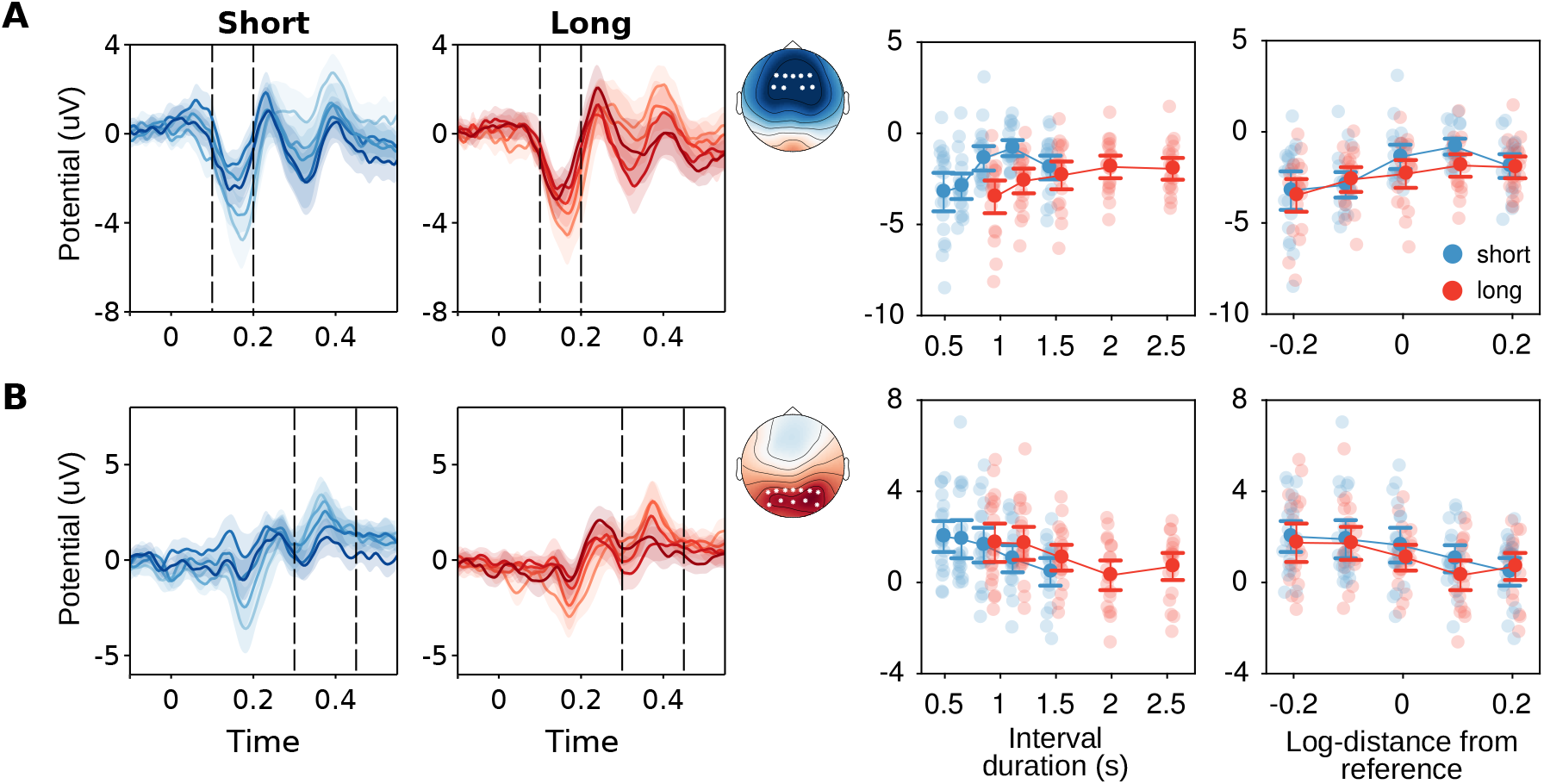
Post-offset ERPs. ERPs evoked at the end of the interval (darker colours corresponding to longer intervals) and reference (blue for the short reference and red for the long reference). Zero indicates the offset of the interval. (A) Activity from early fronto-central electrodes (F3/F1/Fz/F2/F4, FC4/FC2/FCz/FC1/FC3). Dashed lines indicate the period used for the topographies and the error-plots. Topographies represent activity collapsed for the period marked by the dashed line collapsed across intervals and conditions. Right panel shows early fronto-central activity (mean activity ± CI) across conditions as a function of the physical duration of the interval and of its relative distance to the reference (right panel). (B) Activity from late parieto-occipital electrodes (P5/P3/P1/Pz/P2/P4/P6, PO8/PO4/POz/PO3/PO7). Dashed lines indicate the period used for the topographies and the error-plots. Topographies represent activity collapsed for the period marked by the dashed line collapsed across intervals and conditions. Right panel shows late parietooccipital activity (mean activity ± CI) across conditions as a function of the physical duration of the interval and of its relative distance to the reference.

### 2.4 EEG activity and behaviour

In a final step, we investigated whether activity in both the pre-interval and post-interval periods was associated with behavioural performance. An l2-regularised logistic regression was trained on the shortest and longest interval of each reference and was tested on the three central intervals [Gouvêa et al., 2015]. The probability of the interval being predicted as “long” was used as an extra predictor in a logistic regression between the duration of the interval and the probability of the participant responding longer. For the pre-interval offset, there was anecdotal evidence that EEG activity did not improve the prediction of behaviour (one sample t-test, *t*_19_ = 1.20, *p* = 0.2436, *d* = 0.2691, *BF*_10_ = 0.4373). For the post-interval period, there was an association between EEG pattern activity and behaviour (window tested=-50ms-700ms;cluster=280-336, *p* = 0.0306).

## 3 Discussion

In this paper, we used a temporal categorisation task with two temporal references to investigate whether EEG activity encodes time, decision or both. Our results suggest that the two types of information are represented in the pre-offset period, but mainly decision-related information is represented in the post-interval period.

Different theoretical models have been proposed to explain the mechanisms of interval timing [Hass and Durstewitz, 2016]. Several of these models can be grouped into two major categories, depending on the mechanism used to track different durations: models that rely on adaptive threshold settings and models that rely on adaptive clock rates [Balci and Simen, 2016]. In adaptive threshold models, clock signals are generated with a constant rate, and different intervals are timed by accumulating a different number of pulses. In adaptive-rate models, different ranges are estimated by using a different rate of accumulation [Balci and Simen, 2016].

In a previous study, Pfeuty et al. [2005] showed that the CNV slope, but not its amplitude, varies with the reference interval. According to the authors, their results suggest that different rates of pulse accumulation would encode different durations and that the CNV reflects this process. Here, using an extended version of their analysis [Pfeuty et al., 2005], we found a similar pattern of findings. Additionally, we found that the ratio between the slope of the CNV in different blocks approximates the ratio between the two temporal references. Although this finding is compatible with the idea of the CNV reflecting pulse accumulation with different rates, the exact role of the CNV in temporal processing has been largely discussed. Recent studies have shown that decisional and memory mechanisms might also modulate the amplitude of the CNV [Ng et al., 2011; Wiener and Thompson, 2015], indicating that it does not reflect a purely temporal accumulation process. On the other hand, it is worth to point out that: (1) no single event-related component likely reflects a single cognitive function; (2) Decisions and memory mechanisms are an essential aspect on several temporal tasks. Thus, although it is not possible to state that the CNV tracks an accumulator process exclusively, our findings support the idea that the CNV is related to decision-making processes that develop over time and that this process develops at different rates depending on the temporal reference.

In addition to the CNV analysis, we also used MVPA to investigate what information is encoded in the EEG signal. In previous works, we have shown that time-related information was decodable from pre-offset EEG signals in temporal reproduction [Barne et al., 2018] and temporal categorisation tasks [Bueno et al., 2017]. However, because in our previous categorisation task there was only one reference interval [Bueno et al., 2017], time and decision information were intrinsically correlated. In the present study, we show again that it is possible to decode time-related information from within each condition. Critically, we found that pre-offset signals encode information about both time and decision. Because MVPA analysis considers all electrodes, it is possible that temporal information is not contained in or restricted to electrodes where the CNV is found. Importantly, the fact that temporal information is present on the pre-offset EEG is in line with findings that show that behavioural performance for durations longer than the standard, and thus when the CNV has deflected, is still a function of the amount of time that had passed [Kononowicz and van Rijn, 2014].

A similar MVPA analysis performed on the post-interval period showed that EEG activity in this period encoded mainly decision-related information. The time periods of this effect are similar to recent findings that suggest the existence of early potentials related to temporal processing, such as the N1P2 [Kononowicz and van Rijn, 2014] and to several studies that have suggested the existence of a Late Positive Component of timing (LPCt) [Bannier et al., 2019; Gontier et al., 2008; Lindbergh and Kieffaber, 2013; Paul et al., 2003, 2011; Wiener and Thompson, 2015]. However, although several authors have claimed to find an LPCt, this component has been identified relative to different temporal markers and with different topographies. For example, in the study from Wiener and Thompson [2015], the LPCt was measured as the positive potential evoked at frontocentral electrodes after the response was made by the participant and had higher amplitudes for shorter intervals. In other studies, such as [Gontier et al., 2008; Paul et al., 2003, 2011], the LPCt was measured at prefrontal electrodes after the interval offset and were higher for longer intervals. Recently, two studies have reported a positive post-offset potential at centroparietal electrodes, both with higher amplitudes for shorter intervals [Bannier et al., 2019; Lindbergh and Kieffaber, 2013].

In our study, the early post-interval components were found at frontocentral electrodes and the late components at parieto-occipital sites. For the early potential, we found more negative amplitudes as a function of how much shorter the interval was relative to the reference. This pattern is different from what has been reported by [Kononowicz and van Rijn, 2014], that found a u-shaped pattern between time and amplitude. However, the authors used the N1-P2 amplitude, while we focused on the amplitude of the N1. For the LPCt, we found higher amplitudes as a function of how much shorter the interval was relative to the reference, a pattern in agreement with recent findings of Lindbergh and Kieffaber [2013] and Bannier et al. [2019]. For both early and late components, the respective authors suggested that this activity was related to temporal processing [Bannier et al., 2019; Kononowicz and van Rijn, 2014]. However, because they used only one reference interval, it was not possible to dissociate whether the amplitude of these components was related to time itself or to whether the interval was shorter or longer than the reference. On the other hand, our results suggest that the relative distance to the temporal reference strongly and almost exclusively influences the pattern of post-interval brain activity.

Our results are in general agreement with recent proposals that temporal processing can be accomplished based on mechanisms similar to the drift-diffusion model (DDM) of decision-making [Balci and Simen, 2014; Balci and Simen, 2016; Simen et al., 2011]. In this view, temporal categorisation is characterised as a two-stage diffusion process. The first stage corresponds to interval estimation as an evidence accumulation process, while the second stage corresponds to the drift-diffusion model for decision [Balci and Simen, 2014; Balci and Simen, 2016; Simen et al., 2011]. An essential feature of this model during the first stage of interval estimation is the presence of a pattern of activity that scales proportionally to the interval to-be-timed. In non-human animals, this type of evidence comes mainly from ramping cell activity [Durstewitz, 2003; Komura et al., 2001; Reutimann et al., 2004]. In human EEG, it has been proposed that the CNV could represent a similar time adaptive activity. Although this proposal is in agreement with our findings, as mentioned, there are recent criticisms on the relation of the CNV to interval timing. On the other hand, our MVPA results indicated that pre-offset EEG activity encodes not only time but also the relative distance of a given interval to its reference. This result suggests the existence of a time-adaptive activity during this first stage, which is in agreement with the mechanisms of the first diffusion stage.

The second stage starts once the stimulus duration has elapsed and consists of a comparison between the current stimulus-duration and the temporal reference. During this stage, evidence builds towards one of two thresholds, similar to the drift-diffusion model of decision-making. In our results, the pattern of the post-offset activity was modulated mainly by the relative distance of the interval to the reference, consistent with a decisional related signal. This pattern was strongly reflected on the LPCt, a component that has a temporal and scalp distribution that resembles components such as the P300 and the centro-parietal positivity (CPP) which, in turn, have been associated with tracking decision variables of DDM [O’connell et al., 2012; Twomey et al., 2015]. Importantly, the LPCt seems to be stronger for intervals that are shorter than the reference. This is consistent with the proposal that the second stage decision process should take place only for shorter than reference intervals: given that the stimulus duration has passed the long reference duration, a decision can be made with confidence without the second stage [Balci and Simen, 2014; Balci and Simen, 2016; Simen et al., 2011].

In summary, our results suggest that temporal and decisional information is encoded in EEG activity. The dynamics of how these pieces of information are tracked is consistent with recent proposals that approximate temporal processing with decisional models. The findings open new lines of investigation into the possible shared mechanisms of temporal and non-temporal decision making.

## 4 Methods

### 4.1 Participants

Twenty volunteers (age range, 20–32 years; 13 female) gave informed consent to participate. All of them had normal or corrected-to-normal vision and were free from neurological or psychological diseases. The experimental protocol was approved by The Research Ethics Committee of the Federal University of ABC (UFABC) and the experiment was performed in accordance with the approved guidelines and regulations.

### 4.2 Stimuli and Procedure

Stimuli were presented using Psychtoolbox [Brainard, 1997] v.3.0 package for MATLAB on a 17–inch CRT monitor with a vertical refresh rate of 60 Hz, placed 50 cm in front of the participant. Responses were collected via a response box with 9 buttons (DirectIN High Speed Button; Empirisoft). The task consisted of a computerised “shoot the target” task adapted from Guilhardi et al. [2010] and Bueno et al. [2017].

As in Bueno et al. [2017], the task consisted of two types of trials. Regular trials (480 trials, figure 1A) started with the presentation of a target (a bulls-eye with a 1.5 visual degree radius, red and black) at the left hemifield of the screen (background RGB-colour 150;150;150) and a “gun sight” (an empty circle with 0.5 visual degree radius) at the centre of the screen. After a random interval (500 ms–1000 ms) a beep (1000 Hz, 70 dB, 100 ms duration) was presented simultaneously with the start of movement of the target from left to right. A button press (with the right hand) simultaneously produced a “shot” (a blue disc, presented inside the gun sight) and a second beep (500 Hz, 70 dB, 100 ms duration). Participants were instructed to hit the target by pressing the button only once per trial at the appropriate moment.

Test trials (480 trials, figure 1B) followed the same design, but the trajectory of the target was masked by a grey rectangle (3 visual degrees of height, RGB-colour 130;130;130) and an automatic shot was given at one of five possible intervals. At the end of each test trial, participants had to judge whether the shot occurred “before” or “after” the target reached the centre of the screen. Participants responded by pressing two different buttons with their left hand. Responses in test trials were unspeeded and could be given only after the presentation of a response frame at the end of the trial. Trials were presented in a random order, with the restriction of having a maximum of three test trials in sequence.

The target could move at two different speeds (figure 1C): in short trials the target reached gun sight after 0.9 seconds. In long trials, it took the target 1.5 seconds to reach the gun sight. Trials were presented in a blocked design (12 blocks of 80 trials, 40 regular trials and 40 test trials in each block) and the speed of the target varied between blocks (6 blocks for each speed, presented in a random order). The first ten trials of every block were regular trials. In short blocks, the shot in test trials could be presented in one of the following five intervals: 0.54 s, 0.70 s, 0.90 s, 1.16 s and 1.5 s. In long trials the intervals were of 0.90 s, 1.16 s, 1.5 s, 1.94 s and 2.50 s. Test intervals were defined using a logarithmic scale around the references. Each test interval was presented 48 times throughout the experiment. The experimental session lasted 75 minutes on average.

#### 4.2.1 EEG recordings and pre-processing

EEG was recorded continuously from 64 ActiCap Electrodes (Brain Products) at 1000 Hz by a QuickAmp amplifier (Brain Products). All sites were referenced to FCz and grounded to AFz. The electrodes were positioned according to the International 10–10 system. Additional bipolar electrodes registered the electrooculogram (EOG). EEG pre-processing was carried out using BrainVision Analyzer (Brain Products). All data were down-sampled to 250 Hz, re-referenced to the average of the auricular electrodes, filtered (band-pass from 0.05 up to 30 Hz), epoched from 250 ms before the first beep to 1000 ms after the second beep and baselined from −100 ms to 0 ms relative to the first beep. An independent component analysis (ICA) was performed to reject eye movement artefacts. Eye related components were identified by comparing individual ICA components with EOG channels and by visual inspection. The number of trials rejected for each participant was small (4.14% on average).

Event related analysis (ERP) were performed using SPM12 [Litvak et al., 2011], Fieldtrip [Oostenveld et al., 2011] toolboxes and custom written scripts. To analyse post-interval evoked potentials, the same pipeline was used, but with a more aggressive band-pass filter (from 1Hz up to 30Hz, applied at the continuous data), and an epoch from −350 to 750ms relative to the second beep (0.13% of rejected trials on average). For the MVPA analyses, we used 48 electrodes (Fz, F1, F2, F3, F4, F5, F6, F7, F8, FC1, FC2, FC3, FC4, FC5, FC6, Cz, C1, C2, C3, C4, C5, C6, CPz, CP1, CP2, CP3, CP4, CP5, CP6, Pz, P1, P2, P3, P4, P5, P6, P7, P8, POz, PO3, PO4, PO7, PO8, PO9, PO10, Oz, O1, O2); outermost electrodes were excluded from all participants due to a high level of artefacts.

### 4.3 Analysis

#### 4.3.1 Behavioural Analysis

For each participant, two Logistic functions were fitted to the proportion of “after” responses, one for each reference interval. Each function was defined by four parameters: threshold, slope, lapse rate and guess rate[Wichmann and Hill, 2001]. Guess rates and lapse rates were restricted to a maximum of 0.05. Slope and threshold were fitted separately for each participant and condition, using maximum likelihood estimation as implemented in the Palamedes Toolbox [Prins and Kingdom, 2009]. Quality of fit for each participant was assessed with the Tjur’s Coefficient of Determination [Tjur, 2009] (*D* = 0.59 ± 0.02; lowest individual *D* = 0.33).

The points of subjective equality (PSE) were estimated as the predicted interval corresponding to 50% of “after” responses. Temporal sensitivity was assessed by Just Noticeable Difference (JND) scores, which represents half of the absolute difference in seconds between the intervals at which 25% and 75% of “after” responses were given. Weber fractions were estimated as:

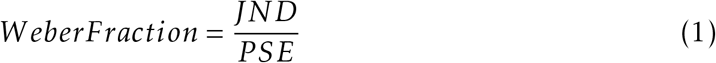

At the group level, the estimated parameters were compared using paired t-tests. When possible, Bayes Factors as implemented in [Krekelberg, 2019] in favour of the alternative hypothesis are reported [Morey and Rouder, 2011; Rouder et al., 2009]. Interpretation of Bayes factors in terms of anecdotal, moderate, strong, very strong and extreme evidence for H0 or H1 was based on the table provided by Wagenmakers et al. [2018]

#### 4.3.2 CNV slope analysis

EEG analysis focused solely on test trials. To estimate the CNV parameters, we used the longest test interval of each reference, as these trials showed the CNV resolution after the reference interval had passed. The CNV, measured in central electrodes (Cz, C1, C2, C3, C4), was considered as the period starting at 300ms after the first beep (thus avoiding its evoked potential) until the presentation of the second beep (1.5 and 2.5s for the short and long reference intervals, respectively). For each participant, CNV data was binned in 100ms windows, and a model combining two linear functions was fitted using a least-square algorithm as implemented in *lsqcurvefit* in MATLAB. The model consisted of three parameters: the moment in time in which the CNV peaked, its maximum amplitude, and its slope. The two linear functions were restricted to have the same slope but with opposite signals. Ratio between the estimated slopes for short and long temporal references was calculated individually by dividing the slope for the short reference by the slope for the long reference. At the group level, the estimated parameters were compared using paired t-tests and Bayes Factors in favour of the alternative hypothesis are reported. The raincloud plot used to show the distribution of the slope ratios was built based on Allen et al. [2019].

#### 4.3.3 Multivariate Pattern Analysis

For the multivariate pattern analysis, we performed pairwise comparisons between each of the ten possible intervals (five possible intervals for the short reference interval and five for the long reference interval). For each comparison, a l2-regularised logistic regression as implemented in *liblinear* [Fan et al., 2008] was trained on part of the data and tested on a separate subset (see below for more details). All model parameters were set to their default values as provided by *liblinear*.

The pre-processing and decoding procedures followed standard guidelines [Lemm et al., 2011]. A leave-one-block-out cross-validation was used to maximise stochastic independence between train and test trials [Lemm et al., 2011]. For each fold, data from 5-blocks were used as the training set and generated predictions to the remaining block. Classification performance for each pairwise comparison was summarised as accuracy and transformed into d-prime.

To investigate how different aspects of the task were represented in EEG activity, two theoretical matrices of pairwise dissimilarity between intervals were built: one based on the physical distances in time and the second based on how different from the reference interval they were (Figure 4, top panels). For ease of comparison, all distances were scaled between 0 and 1. *Within* conditions analysis was performed using upper-left and lower-right quadrants from the theoretical time matrix. These quadrants were used as predictors in a linear regression for the observed classification performance (transformed into d-prime scores) made for each participant and condition (short/long reference interval) separately. In a second analysis, we performed a multiple linear regression between the classification scores observed in the upper-right quadrant with both theoretical matrices as predictors. In both analysis, the estimated linear coefficients were compared, at the group-level, against zero using one-sample t-tests.

For the pre-interval analysis, each logistic regression was fitted to the mean of the last 100ms prior to the end of the intervals, for each participant separately, across the 48 electrodes. For the post-interval EEG activity, pairwise comparisons were done for each time-point. Data from each electrode was smoothed using a moving average of 40 ms before pairwise comparisons were performed. Linear regressions were performed as described above. A cluster-based permutation test (n=10000) was used to correct for multiple comparisons across time, with a clustering threshold of *p* < 0.05 [Maris and Oostenveld, 2007].

#### 4.3.4 Correlation between EEG data and behaviour

In a first step, a l2-regularised logistic regression was trained on the shortest and longest interval of each reference and tested on the three centre intervals. For each test trial, we stored the probability of that trial being classified as long by the classifier. To investigate whether these probabilities were correlated with behaviour, a logistic regression was performed with the physical interval, the condition (short or long block) and the output of the classifier as predictors, and with the actual response of the participant as the dependent variable. To account for the fact that different physical intervals should present a higher probability of being classified as long by the classifier, we normalised these probabilities for each interval based on their mean probability. Data from both conditions (short and long) were collapsed and a single regression was performed for each participant. At the group level, the estimated coefficients were tested using one-sample t-tests.

## 5 Acknowledgements

The authors would like to thank Fuat Balci and the members of the Timing and Cognition Laboratory at UFABC for useful discussions and suggestions on earlier versions of this manuscript. A.M.C. was supported by the São Paulo Research Foundation (FAPESP), research grant #2017/25161-8. V.M.C was supported by grant #2016/19940-1, São Paulo Research Foundation (FAPESP) and Coordination of Superior Level Staff Improvement (CAPES). M.S.C. was supported by by the Brazilian National Research Council (CNPq, grant #465686/2014-1), the Coordination of Superior Level Staff Improvement (CAPES, grant #88887.136407/2017-00), and São Paulo Research Foundation (FAPESP), grant #2014/50909-8. J.R.S. was supported by São Paulo Research Foundation (FAPESP), grant #2018/04654-9 and #2018/21934-5.

